# MicroDLP: Microscope-Enabled Resolution-Switchable 3D Microprinting

**DOI:** 10.64898/2026.10.21.760546

**Authors:** Dimitar Boev, Prajwal Agrawal, Mahmoud Medany, Yong Deng, Gabriele Morlupi, Daniel Ahmed

**Affiliations:** Acoustic Robotics Systems Lab, ARTORG Center for Biomedical Engineering Research, Faculty of Medicine, University of Bern, 3008, Switzerland; Acoustic Robotics Systems Lab, Institute of Robotics and Intelligent Systems, Department of Mechanical and Process Engineering, ETH Zurich, 8092, Switzerland

**Keywords:** additive manufacturing, microscale fabrication, digital light processing, 3D printing, multiscale fabrication

## Abstract

Complex three-dimensional (3D) microarchitectures are increasingly important for microfluidics, microrobotics, organ-on-chip systems, biomedical devices, and architected materials. However, fabricating complex 3D microstructures with micrometre-scale precision remains a major manufacturing challenge. Integrating these fine features with larger supporting structures presents an additional challenge, requiring fabrication across multiple length scales within a single architecture. Although microscale 3D printing enables intricate geometries, the trade-off between resolution and build volume limits multiscale fabrication, while existing approaches often require complex optics and precise registration. Here, we introduce MicroDLP, a microscope-enabled digital light processing platform for versatile, resolution -switchable 3D microprinting. By integrating a projection module into a conventional inverted microscope, MicroDLP exploits its revolving objective turret to switch between 4×, 10×, and 20× objectives, providing projected pixel sizes from 1.89 to 0.378 µm. We characterize photopolymerization, voxel geometry, and lateral feature resolution across magnifications to establish suitable microscale printing conditions. MicroDLP enables the fabrication of diverse and complex 3D microarchitectures, including freestanding trusses with 170 µm struts, microvascular structures with 10 µm features, interlocking gears, suspended microbridges, and microneedles with 3–5 µm tips. Importantly, switching from 10× to 20× during fabrication produces monolithic 3D structures combining larger supporting foundations with fine micropillar arrays, without manual optical realignment. The shared optical pathway also enables coaxial in situ monitoring of photopolymerization. Together, these results establish MicroDLP as a versatile platform for microscale additive manufacturing, combining complex 3D microfabrication with resolution switching within a single monolithic structure.

## 1. Introduction

Fabricating complex three-dimensional (3D) structures at the microscale is central to advances in microfluidics, microrobotics, biomedical devices, and architected materials [1–6]. For many applications, functionality depends on features spanning multiple length scales: microneedles require micrometre-scale tips supported by larger bases for mechanical stability [7, 8]; microfluidic systems incorporate narrow channels, interconnected branches, and suspended elements [9–13]; and microrobotic devices rely on precisely shaped appendages and cavities for actuation and transport [14–21]. These applications highlight two manufacturing challenges: fabricating intricate 3D microarchitectures with high geometric fidelity and integrating different feature scales within a single structure. Conventional photolithography and soft lithography predominantly rely on planar patterning, making complex 3D fabrication challenging and often requiring multiple fabrication and assembly steps [22, 23].

Microscale additive manufacturing (AM) enables direct 3D fabrication but involves trade-offs among resolution, printing speed, and build volume [24–28]. Two-photon polymerization (2PP) achieves submicrometric feature sizes but is constrained by serial voxel writing and often bulky, complex, and expensive optical instrumentation [24]. In contrast, projection microstereolithography (P µSL) and digital light processing (DLP) expose entire layers simultaneously, offering higher fabrication throughput [27, 29]. At fixed optical magnification, increasing the projected area generally enlarges the effective pixel size and limits fine-detail fabrication, whereas higher magnification restricts the printable area [11, 27]. Consequently, a single optical configuration may be insufficient for structures that combine larger supporting elements with intricate microscale features.

Existing multiscale printing approaches address this trade-off through spatially tiled exposures, adjustable magnification, or multiple projection configurations [2, 4, 9, 11, 30, 31]. Advanced P µ SL platforms have demonstrated submicrometre features across centimetre-scale structures using tiled projection [4, 11], while multipath projection stereolithography (MPS) employs separate optical configurations to fabricate large microfluidic devices containing finer features [2]. Commercial multiscale PµSL systems similarly incorporate multiple projection lenses [30]. Although effective, these approaches can introduce additional calibration, optical alignment, and registration requirements. A common optical architecture that enables resolution switching during fabrication could provide a simpler alternative for multiscale 3D microprinting.

A conventional research microscope provides a shared optical architecture for multiscale printing through its revolving objective nosepiece. Its approximately parfocal and coaxial objectives allow magnification, numerical aperture, and field of view to be changed while minimizing optical realignment. Microscope-based projection systems have previously enabled mask-based and maskless micropatterning [10, 22, 23, 32]. However, these approaches focused on two-dimensional fabrication . More recently, Haris et al. integrated DLP into a fluorescence microscope, demonstrating photochemical patterning and proof-of-concept 3D resin printing [33]. However, these studies did not demonstrate objective switching during a single 3D build to integrate different feature scales within a monolithic structure. Extending this capability to 3D printing offers a route to versatile microfabrication and multiscale integration using a common optical architecture with in situ observation.

Mid-print resolution switching presents two technical challenges. First, sufficient geometric registration must be maintained between successive printed layers when objectives are exchanged. Second, changes in magnification, numerical aperture, and irradiance alter photopolymerization behavior and voxel geometry. Consequ ently, characterizing cure depth, lateral resolution, and exposure conditions across objectives is essential for achieving consistent fabrication and structural continuity. Here, we introduce MicroDLP, a microscope-integrated DLP platform for versatile 3D microfabrication and multiscale printing. By coupling a digital micromirror device (DMD) projection module into a conventional inverted microscope, MicroDLP employs interchangeable 4×, 10×, and 20× objectives to adjust fabrication resolution and field of view. We systematically characterize photopolymerization behavior across magnifications and demonstrate the fabrication of diverse complex 3D microarchitectures, including trusses, miniature mechanical structures, microvascular networks, interlocking gears, suspended microbridges, and microneedle arrays with 3–5 µm tips. We further demonstrate mid-print resolution switching from 10× to 20× to integrate fine micropillar arrays onto larger supporting structures within a single monolithic print, without manual optical realignment. The shared microscope optical architecture additionally enables coaxial in situ observation of photopolymerization. Together, these results establish MicroDLP as an accessible and adaptable platform combining versatile 3D microprinting, multiscale integration, and in situ optical monitoring.

## 2. Results

### 2.1. Platform architecture and the parfocal projection path

MicroDLP uses a modular optical and mechanical arrangement to convert a conventional inverted microscope into a versatile 3D microprinting platform (**Fig. 1A**). A printing module containing a resin vat and a vertically translated build platform mounts on the microscope stage, while a DMD-based projection module couples patterned light through a side port (**Fig. 1B**). The selected objective projects each mask onto the resin interface. In addition to enabling microscale manufacture through multiple objectives, this architecture retains a shared observation path to the fabrication region.

**Figure 1.**
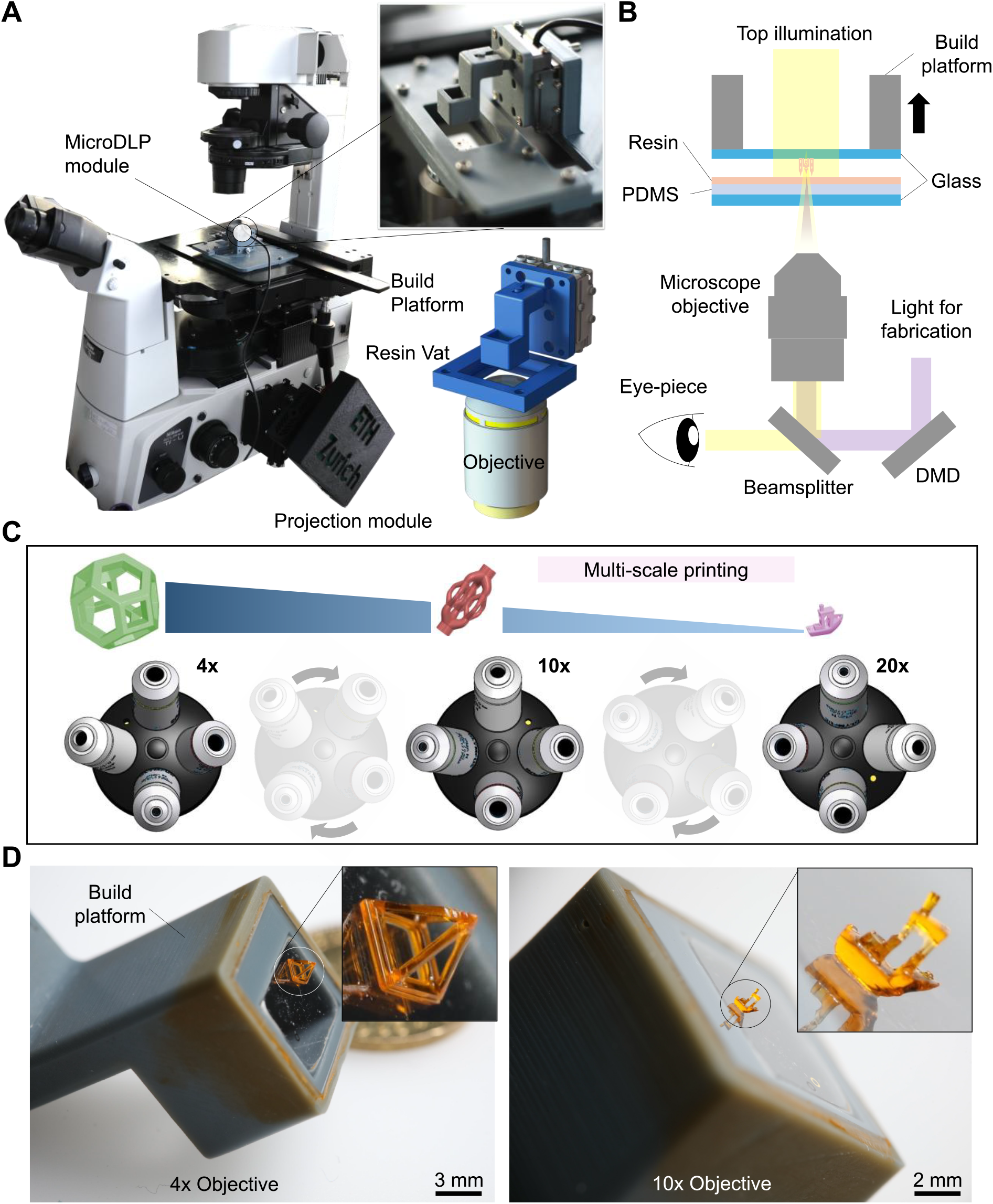
MicroDLP platform and principle of resolution-switchable 3D microprinting. **A.** Overview of the microscope-integrated MicroDLP system, comprising the DMD projection module and the printing module mounted on a standard inverted microscope. The enlarged view shows the detachable build platform and resin vat coupled to the precision Z-stage, enabling rapid removal and exchange of printed structures. **B.** Optical and printing configuration. Patterned light from the DMD is relayed through the selected microscope objective onto the resin interface, while the build platform translates vertically within a vat sealed by a PDMS-coated borosilicate window. A beamsplitter provides a shared optical pathway for projection and in situ observation. **C.** Resolution-switching principle. Rotation of the microscope nosepiece between 4×, 10×, and 20× objectives changes the projected field of view and feature resolution while maintaining approximately the same optical axis and focal plane. **D.** Representative structures fabricated at different magnifications: a pyramidal truss printed at 4× and a Benchy model printed at 10×. Scale bars: 3 mm and 2 mm, respectively. DMD, digital micromirror device; DLP, digital light processing; FOV, field of view; PDMS, polydimethylsiloxane.

In the printing module, a high-precision piezoelectric inertia Z-stage controls the vertical position of a detachable build platform immersed in a vat of photocurable resin, sealed at its base by a borosilicate glass window coated with polydimethylsiloxane (PDMS) (**Figure 1B**). PDMS was selected for two functional reasons. Its low surface energy facilitates the release of polymerized structures from the vat window, reducing adhesion during fabrication [29]. Its high optical transparency across the UV and visible range preserves transmission for both projection and observation [34]. The projection module comprises a commercial DMD evaluation board (DLP6500EVM), a broadband dielectric mirror, and 3D printed optomechanical mounts, which relay patterned images into the microscope optical path through the side port and onto the resin interface through the selected objective.

The key component enabling resolution switching is the microscope’s revolving nosepiece, equipped with 4×, 10×, and 20× objectives (**Figure 1C**). The parfocal and approximately coaxial arrangement of these objectives allows magnification, numerical aperture, field of view, and delivered irradiance to be changed while minimizing shifts in the optical axis and focal plane. In the demonstrated configuration, objective switching was performed without manual optical realignment, enabling fabrication to continue directly on the existing structure (**Figure 4B, C**). This capability enables fabrication resolution to be adjusted during printing, allowing microstructures with different feature scales to be integrated within a single monolithic 3D construct.

**Figure 2.**
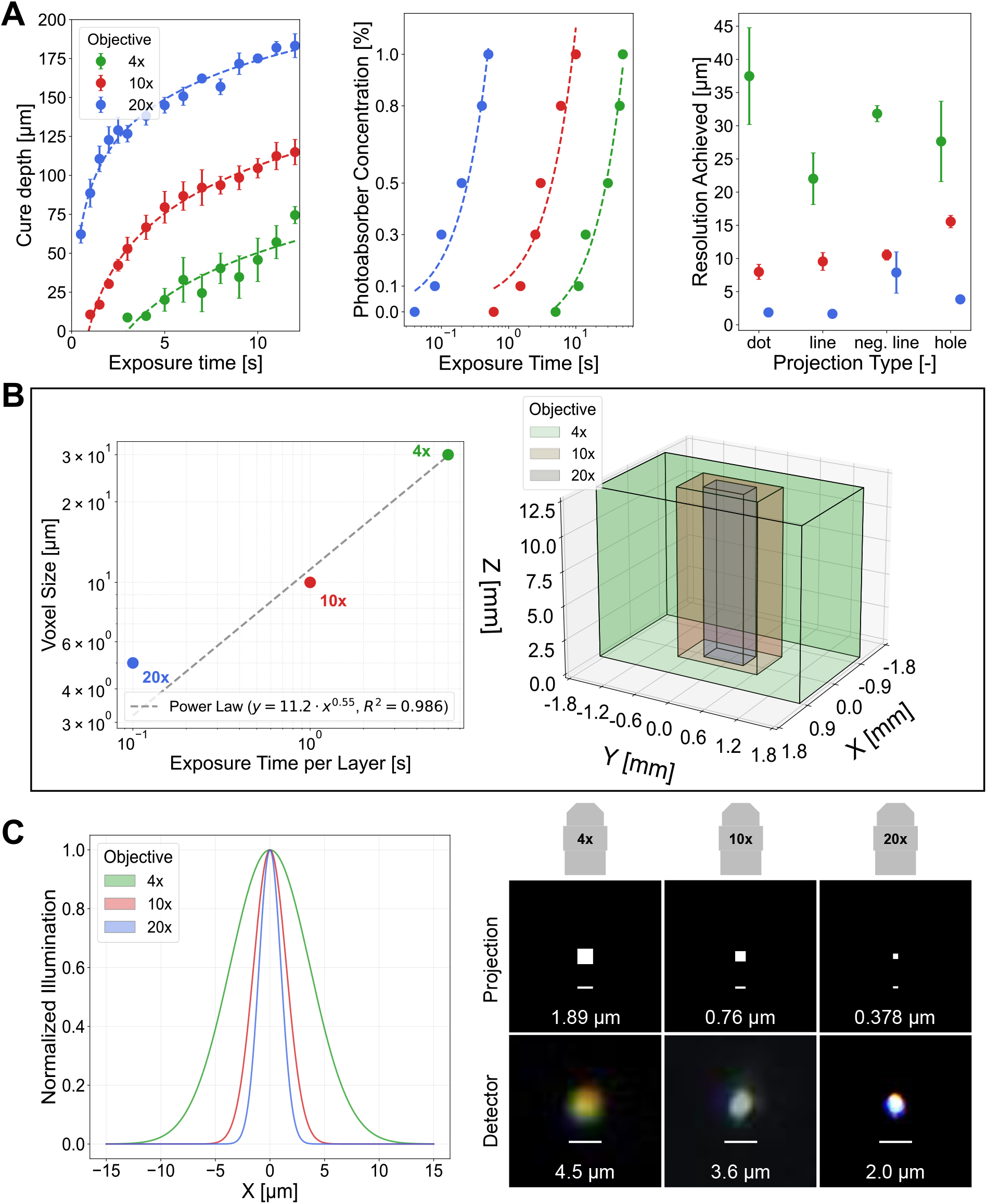
Cross-objective photopolymerization characterization. **A.** Left: cure depth as a function of exposure time for the 4x, 10x, and 20x objectives, showing logarithmic dependence at all magnifications, with the 20x objective reaching greater cure depths at shorter exposures. Center: Sudan I photoabsorber concentration (0 to 1.0 wt%) plotted against the exposure time required to reach the polymerization threshold, shown on a logarithmic time axis, with dashed lines indicating fits. Right: *xy* resolution achieved for dot, line, negative line, and hole projections under each objective, spanning 22 to 37 µm at 4x, 8 to 16 µm at 10x, and 2 to 8 µm at 20x. Data are presented as mean ± s.d. **B.** Left: voxel size as a function of exposure time per layer, fitted to a power law (y = 11.2x^0.55^), yielding 30 µm at 4x, 10 µm at 10x, and 4.8 µm at 20x. Right: maximum build volume accessible under each objective, spanning 2.0 by 3.6 by 12.5 mm in xyz at 4x and decreasing in *xy* with increasing magnification. **C.** Left: simulated normalized single-pixel Gaussian intensity distributions for each objective. Right: projected pixel sizes of 1.89, 0.76, and 0.378 µm for the 4x, 10x, and 20x objectives, and the corresponding spot sizes measured at the detector of 4.5, 3.6, and 2.0 µm.

**Figure 3.**
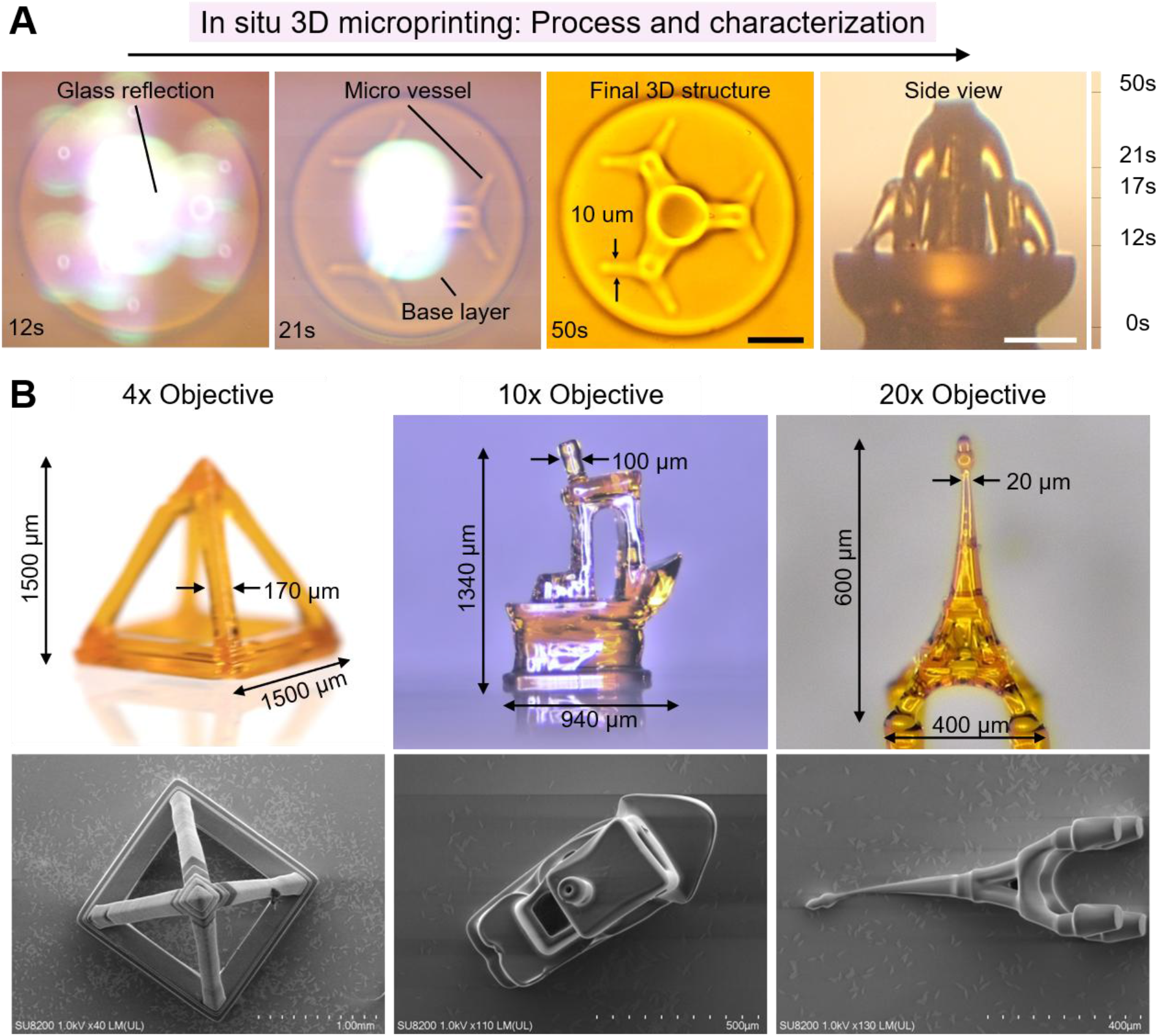
Coaxial in situ monitoring and single-objective fabrication. **A.** In situ monitoring of photopolymerization through the shared optical path during fabrication of a 3D microvessel model. A beamsplitter in the imaging path directs most of the light to the resin and diverts the remainder to a camera, allowing the curing front to be imaged at the numerical aperture used to pattern it. Frames show the initial glass reflection at 12 s, emergence of the base layer and micro vessels at 21 s, and the completed structure carrying 10 µm features at 50 s. The side view shows the accumulated structure over the 0 to 50 s sequence. Scale bars: 30µm. **B.** Structures fabricated at fixed magnification, shown as light micrographs (top) with corresponding SEM images (bottom). The 4x objective produced a tetragonal pyramidal truss 1500 µm in height and 1500 µm across the base, with 170 µm struts of uniform geometry. The 10x objective produced a Benchy boat model measuring 1340 µm by 940 µm with 100 µm topological detail and smooth layer transitions. The 20x objective produced an Eiffel Tower model measuring 600 µm in height and 400 µm across the base, with a minimum feature size of 20 µm. SEM scale bars: 1.00 mm (4x), 500 µm (10x), 400 µm (20x).

**Figure 4.**
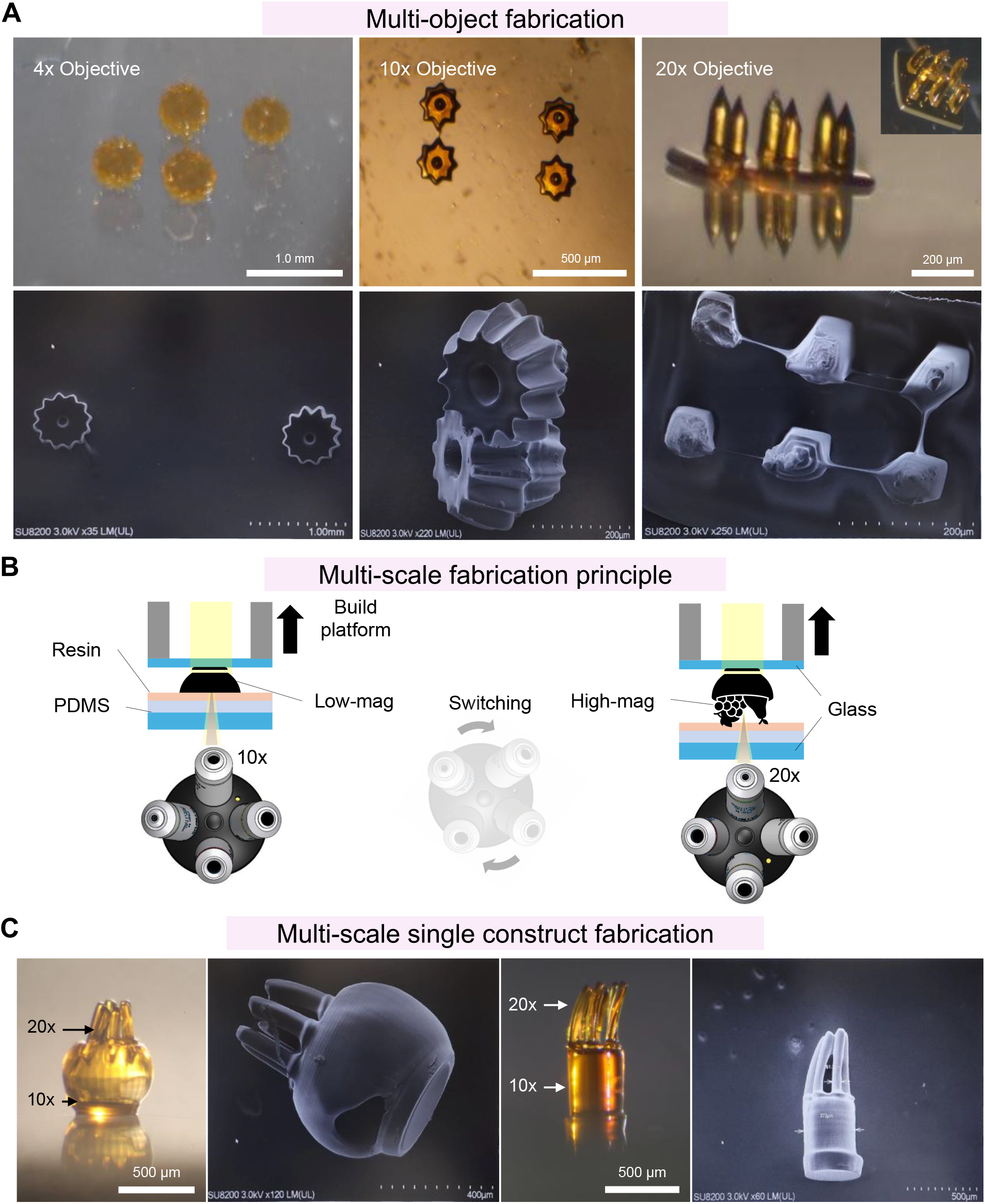
Mid-print magnification switching and multiscale fabrication. **A.** Multi-object fabrication at discrete magnifications within a single build, shown as light micrographs (top) with corresponding SEM images (bottom). The 4x objective produced gears with sharper edges and more clearly defined three-dimensional contours. The 10x objective produced interlocking slanted gears. The 20x objective produced a microneedle array with 200 µm bases and 3 to 5 µm tips, with the inset showing a tilted view, and suspended micro-bridges spanning adjacent supports. SEM scale bars: 1.00 mm (4x), 200 µm (10x), 200 µm (20x). **B.** Schematic of the objective-switching sequence. A low magnification objective (10x) polymerizes the base of the structure against the PDMS-coated glass window while the build platform translates upward, after which the nosepiece is rotated to a high magnification objective (20x) and printing continues on the existing structure without repositioning the platform. **C.** Monolithic constructs produced by switching from 10x to 20x during a single continuous print, shown as light micrographs with corresponding SEM images. Left: a 10x base carrying 20x fine features. Right: a 10x foundation supporting 20x hair-like micropillars. Structural continuity was preserved across the transition in both constructs. SEM scale bars: 400 µm (left), 500 µm (right).

Standard 3D CAD models are sliced using open source or commercial DLP slicers supporting micrometer layer heights, and the resulting image stacks are imported into a custom MicroDLP graphical user interface controlling DMD and Z-stage parameters. During printing the Z-stage executes continuous or quasi-continuous upward motion while the DMD projects a time-sequenced series of binary or grayscale masks, polymerizing resin selectively within illuminated regions.

### 2.2. Cross-objective photopolymerization and resolution scaling

Fabricating complex 3D microstructures at different magnifications requires precise control over photopolymerization, as projected pixel size alone does not determine the achievable feature resolution. Changes in numerical aperture, projected field of view , and delivered irradiance influence curing behavior, voxel geometry, and ultimately the fidelity of printed microstructures.

We therefore characterized exposure-dependent cure depth, polymerization threshold, lateral feature resolution, and voxel geometry using the 4×, 10×, and 20× objectives with a common resin (**Figure 2**). These measurements establish the relationship between optical magnification and achievable microscale fabrication resolution, providing the basis for selecting suitable printing conditions and switching objectives during the fabrication of multiscale 3D structures.

Cure depth increased logarithmically with exposure time across all three objectives, consistent with Jacobs’ working curve (**Figure 2A**). Higher magnification resulted in substantially greater cure depths at shorter exposure times, reflecting differences in the delivered irradiance. For example, at an exposure time of 5s, the 4× objective produced a cure depth of approximately 15 µm, whereas the 20× objective achieved approximately 150 µm, representing a tenfold increase under the same exposure duration. To further control photopolymerization depth, we incorporated Sudan I as a photoabsorber to attenuate light penetration into the resin and confine polymerization along the optical axis. Increasing the Sudan I concentration systematically increased the exposure time required to initiate curing across all magnifications, consistent with reduced light availability for photopolymerization [27, 35, 36]. exposure conditions to be established for microscale and multiscale 3D printing.

To quantify the achievable microscale fabrication resolution, we evaluated a range of projected features, including dots, lines, negative lines, and circular holes (**Figure 2A**). Increasing objective magnification progressively improved lateral resolution, with measured feature sizes ranging from 22–37 µm at 4× to 8–16 µm at 10× and 2–8 µm at 20×, depending on the projected geometry. At higher magnifications, dots and lines exhibited sharper boundaries and improved dimensional fidelity, enabling the fabrication of features at the few-micrometre scale. Circular holes showed greater deviations from the projected geometry, potentially due to local exposure variations, oxygen inhibition, and polymerization-induced shrinkage. The 4× objective provided a larger fabrication area but exhibited more pronounced edge broadening, particularly for narrow lines and holes. These results highlight the trade-off between fabrication area and spatial resolution, demonstrating how objective magnification can be selected to fabricate complex 3D microstructures with different feature sizes and geometries.

We further investigated the relationship between exposure time and voxel dimensions, revealing a nonlinear increase in voxel size that was well described by a power-law relationship (**Figure 2B**). The accessible fabrication area decreased with increasing objective magnification, with the 4× objective providing a projected area of approximately 2.0 × 3.6 mm², whereas the 20× objective enabled finer features within a smaller field of view. To examine the optical basis of this resolution scaling, we simulated single-pixel Gaussian intensity profiles, which showed progressively narrower illumination spots at higher magnifications (**Figure 2C**). The calculated projected pixel sizes were 1.89, 0.76, and 0.378 µm for the 4×, 10×, and 20× objectives, respectively, corresponding to measured optical spot sizes of 4.5, 3.6, and 2.0 µm . These measurements demonstrate how optical magnification influences the illumination distribution and, consequently, the achievable feature dimensions. Under the investigated exposure conditions, higher magnifications produced smaller lateral voxel dimensions but greater axial elongation, highlighting the importance of controlling polymerization in both lateral and vertical directions. By adjusting exposure time and layer height, we established suitable printing conditions for each objective to balance feature resolution, structural continuity, and fabrication throughput. Together, these results provide the experimental basis for fabricating complex 3D microarchitectu res at different resolutions and integrating multiple feature scales within a single monolithic structure.

### 2.3. Coaxial *in situ* monitoring of the curing front

An additional advantage of microscope-enabled 3D microfabrication is the ability to observe the curing region through the same objective used to project the printing patterns (**Figure 3A**). A beamsplitter directs a fraction of the optical signal to a camera, enabling in situ visualization of photopolymerization and the developing microstructure. This coaxial configuration provides direct visual monitoring of the fabrication process without interrupting printing, although automated feedback control has not yet been implemented. Time-lapse imaging during fabrication of a 3D microvascular structure demonstrates this capability (**Figure 3A**). Following the initial reflection from the glass interface at 12 s, polymerized microvascular features became visible at approximately 21 s, and the completed structure, containing features as small as 10 µm, was observed at 50 s. These observations demonstrate the ability to visualize the progressive formation of complex 3D microstructures in real time, providing a foundation for future closed-loop monitoring and correction of fabrication deviations.

### 2.4. Versatile 3D microprinting of complex microstructures

To demonstrate the versatility of MicroDLP for fabricating complex three-dimensional microstructures, we printed representative geometries at different objective magnifications and characterized them using optical microscopy and scanning electron microscopy (SEM) (**Figure 3B**). The 4× objective produced an open pyramidal truss approximately 1500 µm in height, with struts measuring 170 µm. At 10×, we fabricated a miniature Benchy boat measuring approximately 1340 × 940 µm, incorporating curved surfaces and geometric details down to 100 µm . The 20× objective enabled the fabrication of an Eiffel Tower structure measuring approximately 600 × 400 µm, with fine structural elements as small as 20 µm. Together, these examples demonstrate the ability to fabricate diverse 3D microarchitectures incorporating freestanding elements, intricate geometries, curved surfaces, and microscale features. Each structure was fabricated using the exposure conditions established for its respective objective (Section 2.2). Optical and SEM characterization confirmed the formation of well-defined 3D geometries, with continuous structural features and no apparent delamination. These results demonstrate the versatility of MicroDLP for fabricating complex microstructures across different length scales while highlighting the trade-off between fabrication area and achievable feature resolution. This limitation motivates switching objective magnification during fabrication to integrate larger structural features and finer microscale details within a single monolithic 3D structure.

### 2.5. Resolution-switchable 3D microprinting of complex microarchitectures

We next investigated two complementary fabrication modes enabled by objective switching (**Figure 4**). In the first mode, distinct 3D microstructures were fabricated at different magnifications on a shared substrate, demonstrating the geometric versatility of MicroDLP. In the second mode, objective magnification was changed during fabrication of a single structure, enabling larger supporting geometries and finer microscale features to be integrated into a monolithic construct. Together, these modes demonstrate the versatility of microscale 3D fabrication and the capability for resolution -switchable multiscale printing.

In multi-object mode, MicroDLP fabricated gears at 4×, three-dimensional interlocking slanted gears at 10×, and microneedle arrays and suspended microbridges at 20× (**Figure 4A**). The microneedles featured bases of approximately 200 µm tapering to tips as small as 3–5 µm . The interlocking gears and suspended microbridges further demonstrate the ability to fabricate complex 3D geometries, including interacting structural elements and freestanding features. These results highlight the versatility of MicroDLP in producing diverse microarchitectures with different geometric complexities and feature dimensions within a single fabrication session.

To demonstrate monolithic multiscale fabrication, a supporting foundation was first printed using the 10× objective, after which printing was paused and the revolving nosepiece was rotated to the 20× objective (**Figure 4B**). During this transition, the Z-stage maintained its position, and the partially fabricated structure remained attached to the build platform and submerged in resin. Exposure time and layer height were adjusted to the operating conditions established for the 20× objective (Section 2.2), allowing high-resolution features to be fabricated directly onto the existing foundation without manual optical realignment or repositioning of the structure.

Optical and SEM imaging confirmed the successful fabrication of monolithic structures combining features produced at different magnifications (**Figure 4C**). In one construct, a larger foundation printed at 10× supported an array of fine micropillars fabricated at 20×, while a second structure incorporated hair-like micropillars extending from a larger supporting base. The observed structural continuity across the interfaces demonstrates the feasibility of integrating distinct feature scales within a single 3D microarchitecture. Although quantitative registration accuracy and interfacial mechanical strength remain to be established, these results demonstrate the central capability of MicroDLP: resolution-switchable fabrication of complex, monolithic 3D microstructures at the microscale.

## 3. Discussion

MicroDLP demonstrates a versatile approach to microscale 3D printing by repurposing the revolving objective turret of a conventional microscope as an active scale-switching mechanism. By exploiting the parfocal and approximately coaxial arrangement of microscope objectives, the platform enables changes in fabrication resolution and field of view without requiring multiple dedicated projection pathways. We characterized photopolymerization across 4×, 10×, and 20× objectives using a common resin, obtaining projected pixel sizes of 1.89, 0.76, and 0.378 µm, respectively, and measured optical spot sizes of 4.5 µm at 4× and 2.0 µm at 20×. The platform enabled the fabrication of diverse complex 3D microarchitectures, including pyramidal trusses, miniature mechanical structures, microvascular networks, interlocking gears, suspended microbridges, and microneedle arrays with 3–5 µm tips. Beyond fabricating individual microstructures, switching from 10× to 20× during a single build enabled larger supporting structures to be integrated with fine micropillar arrays within a monolithic print, with structural continuity observed across the interface. These results demonstrate both versatile 3D microfabrication and the ability to adjust fabrication scale during printing. Furthermore, the shared optical architecture enables coaxial in situ observation of photopolymerization, as demonstrated through time-lapse imaging of microvascular structures. Together, these capabilities establish MicroDLP as a platform combining complex 3D microprinting, multiscale integration, and optical process monitoring within a conventional microscope architecture.

Several factors currently limit the platform. Objective switching is performed manually, introducing pauses during fabrication and limiting the frequency of magnification changes. Motorizing the nosepiece and synchronizing objective rotation with the printing sequence could enable automated resolution switching. The lateral printing area is constrained by the field of view of the selected objective, although automated stage translation and spatially tiled exposures could extend the accessible fabrication area. Achievable printing resolution depends on projected pixel size, optical spot dimensions, exposure conditions, and resin photochemistry. Although the projected pixel size reaches 0.378 µm at 20×, experimentally demonstrated features remain in the micrometre range. Further optimization of the optical configuration, photoabsorber concentration, and exposure conditions may improve resolution and geometric fidelity. Curved features exhibited reduced fidelity relative to dots and lines, potentially reflecting local exposure variations and polymerization-induced shrinkage. Grayscale dose modulation may improve geometric accuracy, while integrating coaxial imaging with exposure and focus control could enable closed-loop photopolymerization. Quantitative evaluation of objective-switching repeatability and registration accuracy would further establish the precision limits of multiscale fabrication.

MicroDLP offers opportunities for fabricating complex and hierarchical 3D microarchitectures across multiple length scales. Microneedles combining fine tips with larger supporting bases could support transdermal drug delivery and biosensing, while interconnected 3D microfluidic networks could advance organ -on-chip systems and tissue engineering. Similarly, integrating compliant elements, fluidic channels, and functional microstructures within larger robotic bodies could support the development of microrobotics and biohybrid systems. Additional applications include micro-optics, mechanical metamaterials, and architected soft materials. Integration with biocompatible resins and controlled fabrication environments could further extend the platform toward biofabrication, although compatibility with living cells and biological materials requires further investigation. The underlying approach may also be adapted to other research microscopes with compatible optical configurations. By combining versatile 3D microfabrication with resolution switching during printing, MicroDLP provides an adaptable approach to manufacturing complex microscale structures and integrating different feature scales within monolithic architectures.

## 4. Materials and Methods

### MicroDLP Printer Setup

The MicroDLP printer comprised two modules, a printing module and a digital micromirror device (DMD) projection module, both mounted on an inverted Nikon Ti-U microscope (Nikon, Japan).

The printing module consisted of a translational piezoelectric inertia drive Z stage (PD1/M, Thorlabs, USA) and a resin vat sealed at its base by a borosilicate glass window coated with polydimethylsiloxane (PDMS), which served as the delamination layer. PDMS (SYLGARD 184, Sigma Aldrich, USA) was mixed with curing agent at a 10:1 ratio, degassed under vacuum, spin coated onto the window at 700 rpm for 1 min, and cured at 85 °C for 2 h. Custom components were designed in SolidWorks (Dassault Systèmes, France) and fabricated from Aqua-Grey 8K resin on a Sonic Mini 8K printer (Phrozen Tech, Taiwan), comprising a Z stage adapter plate, a detachable base plate, the resin vat, and an adapter base that mounted the assembly to the microscope stage with four M3 bolts. A borosilicate glass coverslip serving as the fabrication surface was bonded to the detachable base plate using two-component adhesive (UHU 2 min, UHU, Germany).

The projection module consisted of a DLP6500EVM evaluation board (Texas Instruments, USA), a cooling fan, a broadband dielectric mirror (BB07-E02, Thorlabs, USA), and custom optomechanical mounts printed from polylactic acid filament on an Ender 3 S1 printer (Creality, China). Patterned images were relayed into the microscope optical path through the side port and projected onto the resin interface through the selected objective. An 80/20 beamsplitter (Nikon, Japan) directed 80% of the light to the resin and 20% to a Canon EOS camera (Canon, Japan) mounted at the microscope eyepiece, enabling in situ monitoring of photopolymerization through the same optical path used for patterning.

### Materials

Poly (ethylene glycol) diacrylate (PEGDA, average Mn 700 and Mn 250), Sudan I photoabsorber, 2-hydroxy-2-methylpropiophenone photoinitiator (97%), trimethylolpropane triacrylate (TMPTA), Isopropanol (IPA), and the SYLGARD 184 silicone elastomer kit were purchased from Sigma Aldrich, USA. All materials were used as received without further purification.

### Resin Preparation

Resins were prepared by combining PEGDA and TMPTA with 2-hydroxy-2-methylpropiophenone as photoinitiator and Sudan I as photoabsorber. Components were mixed until homogeneous and stored in light-protected containers until use. Photoabsorber concentration was varied from 0 to 1.0 wt% for the cure characterization experiments.

### Printing Procedure

Three-dimensional models were sliced into image stacks (using open-source software slicers like Chitubox or Slic3r) at micrometer layer heights and imported into the MicroDLP control interface. During fabrication, the Z stage executed continuous upward translation while the DMD projected the corresponding sequence of binary or grayscale masks, polymerizing resin selectively within illuminated regions. For multiscale prints, the objective nosepiece was rotated at the designated point in the sequence and exposure time and layer height were switched to the operating point established for the incoming objective, after which printing continued on the existing structure. Printed structures were removed from the build platform, washed with isopropanol (IPA), and dried before characterization.

### Software

A custom graphical user interface was developed in Python using the PyQt5 framework. The interface-controlled projection of sliced model images, Z stage translation, and projection parameters including exposure time, grayscale level, and xy offset, all adjustable in real time during fabrication.

### Characterization

Cure depth was measured as a function of exposure time for each objective at fixed photoabsorber concentration, and the exposure required to reach the polymerization threshold was determined across the photoabsorber range. Resolution in xy was evaluated using projected dot, line, negative line, and hole features under each objective. Voxel volume was measured as a function of exposure time per layer and fitted to a power law. Single-pixel Gaussian intensity distributions were simulated for each objective and compared against intensity profiles measured by camera at the resin plane. Printed structures were imaged by light microscopy and by scanning electron microscopy (SU8200, Hitachi High-Technologies, Japan).

## Author contributions

D.A. conceived and supervised the project. D.B., P.A., and G.M. developed the methodology. D.B. and P.A. conducted the experimental investigations. D.B., P.A., G.M., Y.D., M.M., and D.A. contributed to writing, reviewing, and editing the manuscript. All authors read and approved the final manuscript.

## Funding

D.A. acknowledges funding from the European Research Council (ERC) under the European Union’s Horizon 2020 Research and Innovation Programme (Grant Agreement No. 853309, SONOBOTS), the European Union’s Horizon Europe Research and Innovation Programme and the Swiss State Secretariat for Education, Research and Innovation (SERI) under the SONOCRAFT project (Grant Agreement No. 101187842), the Swiss National Science Foundation (SNSF; MINT 2022, Grant Agreement No. 213058), and the European Research Council under the European Union’s Horizon Europe Research and Innovation Programme through the ERC Proof of Concept project UniADLP (Grant Agreement No. 101248750). The authors also acknowledge financial support from Innosuisse, the Swiss Innovation Agency, under the project “SonoPrint – Ultrasound-Assisted Multimaterial Bioprinter” (Grant No. 129.628 IP-ENG). P.A. acknowledges support from Innosuisse through the BRIDGE Proof of Concept programme (Grant Agreement No. 218795) and the Innovation Booster Robotics programme.

## Data Availability Statement

The data supporting the findings of this study are presented in the article and its Supporting Information. Additional data are available from the corresponding author upon reasonable request.

## Competing interests

The authors declare no competing interests.

## Notes

### Competing Interest Statement

The authors have declared no competing interest.

## References

[1] A. V. Borovjagin, Ogle, B. M., Berry, J. L., & Zhang, J., From Microscale Devices to 3D Printing, Circulation Research 120 (2017) 150–165, 10.1161/CIRCRESAHA.116.308538

[2] Z. J. Geffert, Xiong, Z., Grutzmacher, J., Wilderman, M., Mohammadi, A., Filip, A., Li, Z., & Soman, P., Multipath Projection Stereolithography for Three-Dimensional Printing Microfluidic Devices, ACS Applied Materials & Interfaces 16 (2024) 69807–69817, 10.1021/acsami.4c10547

[3] G. Vunjak-Novakovic, Ronaldson-Bouchard, K., & Radisic, M., Organs-on-a-chip models for biological research, Cell 184 (2021) 4597–4611, 10.1016/j.cell.2021.08.005

[4] X. Zheng, Smith, W., Jackson, J., Moran, B., Cui, H., Chen, D., Ye, J., Fang, N., Rodriguez, N., Weisgraber, T., & Spadaccini, C. M., Multiscale metallic metamaterials, Nature Materials 15 (2016) 1100–1106, 10.1038/nmat4694

[5] Y. Wang, Sun, C., & Ahmed, D., A smart acoustic textile for health monitoring, Nature Electronics 8 (2025) 485–495, 10.1038/s41928-025-01386-2

[6] M. Medany, Mukkavilli, S. K., & Ahmed, D., AI-driven autonomous microrobots for targeted medicine, Nature Reviews Bioengineering 2 (2024) 914–915, 10.1038/s44222-024-00232-y

[7] J. W. Lee, Park, J.-H., & Prausnitz, M. R., Dissolving microneedles for transdermal drug delivery, Biomaterials 29 (2008) 2113–2124, 10.1016/j.biomaterials.2007.12.048

[8] J. Durrer, Agrawal, P., Ozgul, A., Neuhauss, S. C. F., Nama, N., & Ahmed, D., A robot-assisted acoustofluidic end effector, Nature Communications 13 (2022) 6370, 10.1038/s41467-022-34167-y

[9] E. Behroodi, Latifi, H., Bagheri, Z., Ermis, E., Roshani, S., & Salehi Moghaddam, M., A combined 3D printing/CNC micro-milling method to fabricate a large-scale microfluidic device with the small size 3D architectures: an application for tumor spheroid production, Scientific Reports 10 (2020) 22171, 10.1038/s41598-020-79015-5

[10] R. Paul, Zhao, Y., Coster, D., Qin, X., Islam, K., Wu, Y., & Liu, Y., Rapid prototyping of high-resolution large format microfluidic device through maskless image guided in-situ photopolymerization, Nature Communications 14 (2023) 4520, 10.1038/s41467-023-40119-x

[11] C. Vidler, Crozier, K., & Collins, D., Ultra-resolution scalable microprinting, Microsystems & Nanoengineering 9 (2023) 67, 10.1038/s41378-023-00537-9

[12] M. Medany, Piglia, L., Achenbach, L., Mukkavilli, S. K., & Ahmed, D., Model-based reinforcement learning for ultrasound-driven autonomous microrobots, Nature Machine Intelligence 7 (2025) 1076–1090, 10.1038/s42256-025-01054-2

[13] M. Medany, Nama, N., & Ahmed, D., Ultrasound-induced Particle Dynamics in Pathological Vascular Vortices, BioRxiv (2025) 2025.2010.2023.684129, 10.1101/2025.10.23.684129

[14] D. Ahmed, Sukhov, A., Hauri, D., Rodrigue, D., Maranta, G., Harting, J., & Nelson, B. J., Bioinspired acousto-magnetic microswarm robots with upstream motility, Nature Machine Intelligence 3 (2021) 116–124, 10.1038/s42256-020-00275-x

[15] D. Ahmed, Ozcelik, A., Bojanala, N., Nama, N., Upadhyay, A., Chen, Y., Hanna-Rose, W., & Huang, T. J., Rotational manipulation of single cells and organisms using acoustic waves, Nature Communications 7 (2016) 11085, 10.1038/ncomms11085

[16] S. Zhao, Tian, Z., Shen, C., Yang, S., Xia, J., Li, T., Xie, Z., Zhang, P., Lee, L. P., Cummer, S. A., & Huang, T. J., Topological acoustofluidics, Nature Materials 24 (2025) 707–715, 10.1038/s41563-025-02169-y

[17] P. Amado, Dillinger, C., Bahou, C., Hashemi Gheinani, A., Obrist, D., Burkhard, F., Ahmed, D., & Clavica, F., Ultrasound-activated cilia for biofilm control in indwelling medical devices, Proceedings of the National Academy of Sciences 122 (2025) e2418938122, doi:10.1073/pnas.2418938122

[18] S. Yang, Rufo, J., Chen, Y., Chen, C., Drinkwater, B. W., Lee, L. P., & Huang, T. J., Acoustic tweezers for advancing precision biology and medicine, Nature Reviews Methods Primers 5 (2025) 49, 10.1038/s43586-025-00415-w

[19] A. Del Campo Fonseca, & Ahmed, D., Ultrasound Robotics for Precision Therapy, Advanced Drug Delivery Reviews 205 (2024) 115164, 10.1016/j.addr.2023.115164

[20] Z. Shi, Zhang, Z., Schnermann, J., Neuhauss, S. C. F., Nama, N., Wittkowski, R., & Ahmed, D., Ultrasound-driven programmable artificial muscles, Nature 646 (2025) 1096–1104, 10.1038/s41586-025-09650-3

[21] Y. Deng, Paskert, A., Zhang, Z., Wittkowski, R., & Ahmed, D., An acoustically controlled helical microrobot, Science Advances 9 (2023) eadh5260, 10.1126/sciadv.adh5260

[22] J. C. Love, Wolfe, D. B., Jacobs, H. O., & Whitesides, G. M., Microscope Projection Photolithography for Rapid Prototyping of Masters with Micron-Scale Features for Use in Soft Lithography, Langmuir 17 (2001) 6005–6012, 10.1021/la010655t

[23] D. W. Palmer, & Decker, S. K., Microscopic circuit fabrication on refractory superconducting films, Review of Scientific Instruments 44 (1973) 1621–1624, 10.1063/1.1686013

[24] S. O’Halloran, Pandit, A., Heise, A., & Kellett, A., Two-Photon Polymerization: Fundamentals, Materials, and Chemical Modification Strategies, Advanced Science 10 (2023) 2204072, 10.1002/advs.202204072

[25] M. A. S. R. Saadi, Maguire, A., Pottackal, N. T., Thakur, M. S. H., Ikram, M. M., Hart, A. J., Ajayan, P. M., & Rahman, M. M., Direct Ink Writing: A 3D Printing Technology for Diverse Materials, Advanced Materials 34 (2022) 2108855, 10.1002/adma.202108855

[26] S. Subedi, Liu, S., Wang, W., Naser Shovon, S. M. A., Chen, X., & Ware, H. O. T., Multi-material vat photopolymerization 3D printing: a review of mechanisms and applications, npj Advanced Manufacturing 1 (2024) 9, 10.1038/s44334-024-00005-w

[27] C. Sun, Fang, N., Wu, D. M., & Zhang, X., Projection micro-stereolithography using digital micro-mirror dynamic mask, Sensors and Actuators A: Physical 121 (2005) 113–120, 10.1016/j.sna.2004.12.011

[28] J. T. Toombs, Luitz, M., Cook, C. C., Jenne, S., Li, C. C., Rapp, B. E., Kotz-Helmer, F., & Taylor, H. K., Volumetric additive manufacturing of silica glass with microscale computed axial lithography, Science 376 (2022) 308–312, 10.1126/science.abm6459

[29] D. Han, Yang, C., Fang, N. X., & Lee, H., Rapid multi-material 3D printing with projection micro-stereolithography using dynamic fluidic control, Additive Manufacturing 27 (2019) 606–615, 10.1016/j.addma.2019.03.031

[30] C. Xia, & Xu, J., A multi-scale system for projection micro stereolithography, United States Patent No. US20230398739A1. (2023), https://patents.google.com/patent/US20230398739A1/en

[31] P. Chansoria, Winkelbauer, M., Zhang, S., Janiak, J., Liu, H., Boev, D., Morandi, A., Grange, R., & Zenobi-Wong, M., Structured Light Projection Using Image Guide Fibers for In Situ Photo-biofabrication, Advanced Materials 37 (2025) 2419350, 10.1002/adma.202419350

[32] S. Singh-Gasson, Green, R. D., Yue, Y., Nelson, C., Blattner, F., Sussman, M. R., & Cerrina, F., Maskless fabrication of light-directed oligonucleotide microarrays using a digital micromirror array, Nature Biotechnology 17 (1999) 974–978, 10.1038/13664

[33] U. Haris, Plank, J. T., Li, B., Page, Z. A., & Lippert, A. R., Visible Light Chemical Micropatterning Using a Digital Light Processing Fluorescence Microscope, ACS Central Science 8 (2022) 67–76, 10.1021/acscentsci.1c01234

[34] F. Schneider, Draheim, J., Kamberger, R., & Wallrabe, U., Process and material properties of polydimethylsiloxane (PDMS) for Optical MEMS, Sensors and Actuators A: Physical 151 (2009) 95–99, 10.1016/j.sna.2009.01.026

[35] E. Fleck, Sunshine, A., DeNatale, E., Keck, C., McCann, A., & Potkay, J., Advancing 3D-Printed Microfluidics: Characterization of a Gas-Permeable, High-Resolution PDMS Resin for Stereolithography, Micromachines 12 (2021) 1266, 10.3390/mi12101266

[36] M. P. Lee, Cooper, G. J. T., Hinkley, T., Gibson, G. M., Padgett, M. J., & Cronin, L., Development of a 3D printer using scanning projection stereolithography, Scientific Reports 5 (2015) 9875, 10.1038/srep09875

